# Determination of glucosinolate contents in *Brassica* germplasm collections and inter- & intra-leaves distribution pattern using UPLC-MS/MS Multiple Reaction Monitoring scan mode

**DOI:** 10.1101/569889

**Authors:** Awraris Derbie Assefa, Susanna Choi, Jae-Eun Lee, Jung-Sook Sung, On-Sook Hur, Na-Young Ro, Ho-Sun Lee, Jaejong Noh, Aejin Hwang, Ho-Cheol Ko, Yun-Jo Chung, Ju-Hee Rhee

## Abstract

Intact glucosinolate (GSL) profile (five aliphatic; three aromatic, and one indolic glucosinolate) in the leaves of 50 germplasm collections and commercial cultivars of *Brassica rapa*, *Brassica juncea*, and *Brassica oleracea* collected from six different countries and grown under uniform cultural conditions were compared by UPLC-MS/MS. Total GSLs content ranged from 36.80 to 2383.12 μmol/kg DW. Aliphatic GSLs predominated among the entire samples representing from 23.0 to 98.9% of the total GSLs content, where gluconapin and glucobrassicanapin contributed the greatest proportion. Other GSLs such as, progoitrin (PRO), glucotropaeolin (TRO), and glucobarbarin (BAR) were found in relatively low concentrations. Principal Component Analysis (PCA) yielded three principal components with eigenvalue ≥ 1, representing 70.33% of the total variation across the entire data set. Accessions IT260822 & IT32750, and commercial cultivar, “Hangamssam2”, were well distinguished from other samples in the PCA plot due to their significantly high amount of BAR, glucobrassicin (GBC), and glucoerucin (ERU), respectively. The inter- and intra-leaf variations of GSLs were examined in three kimichi cabbage varieties. The GSLs content varied significantly among leaves in different positions (outer, middle, and inner) and sections within the leaves (top, middle, bottom, green/red, and white). Higher GSL contents were observed in the proximal half & white section of the leaves and inner positions (younger leaves) in most of the samples. GBC, gluconasturtiin (NAS), and glucoberteroin (BER) should be studied profusely in *Brassica* plants as some of their degradation products of GBC and NAS are useful in cancer chemopreventive functions, whereas BER takes part in the process of suppressing aging of the skin. GSLs are regarded as allelochemicals; hence, the data related to the patterns of GSLs within the leaf and between leaves at different position could be useful to understand the defense mechanism of *Brassica* plants. The observed variability could be useful for breeders to develop *Brassica* crops with high GSL content or specific profiles of GSLs as required.

## Introduction

Glucosinolates (GSLs) also called β-thioglucoside-N-hydroxysulfates are class of sulfur-containing important plant secondary metabolites naturally occurring in almost all *Brassica* species [1]. GSLs could be classified as aliphatic, aromatic, and indolic glucosinolates based on their side chain structure (R group) which are derived from the amino acid precursors methionine (but also alanine, leucine, isoleucine, or valine in some cases), phenylalanine, and tryptophan, respectively [2]. Most glucosinolates share a basic chemical structure consisting of a β-D-glucopyranose residue linked via a sulfur atom to a (*Z*)-N-hydroximinosulfate ester and a variable R group [3]. Upon hydrolysis by myrosinases, glucosinolates produce several bioactive products including, isothiocyanates, thiocyanates, and nitriles. Glucosinolates and their biosynthetic products are implicated to reduce the risk of cancer in human [4, 5]. Reports also show that breakdown products of GSLs displayed antimicrobial activity [6, 7]. Although their contribution is complex to understand, GSLs are regarded as an important component of flavor in cooked vegetables [8]. GSLs and/or their degradation products serve as a feeding deterrent to wide range of herbivores such as birds, mammals, mollusks, aquatic invertebrates, nematodes, bacteria, and fungi [8, 9]. On the other hand, they also serve to attract and stimulate specialist herbivores such as the larvae of the lepidopteran species *Plutella xylostella* and *Pieris rapae* [9]. The biocidal activity of glucosinolate containing *Brassica* plants made them as a promising alternative to synthetic pesticides for pest and disease control [10]. *In planta* studies of various Brassicaceae seedlings have also showed a positive correlation between specific and/or total GLS contents and disease severity [11].

Glucosinolates are found in the vegetative and reproductive tissues of various dicotyledonous plant families, and are the major secondary metabolites in mustard-oil plants of *Brassicaceae* family [3, 12]. The content of glucosinolate accounts about 1% of the dry weight in *Brassica* vegetables and can go up to 10% in seeds of some plants [3]. The qualitative and quantitative profiles of total and individual GSLs in Brassica vegetables varies significantly due to several factors such as cultivar genotype [13, 14], developmental stage [15], environmental conditions (temperature, light, water, and soil) [16–19], growing seasons [20], agricultural practices [21], level of insect damage [19, 22], and post-harvest conditions [23]. A wide geographic and evolutionary variation is recorded in broccoli [20], *Arabidopsis thaliana* [24], Chinese cabbage [14], and cabbage (*Brassica oleracea* L.) [25]. Apart from the aforementioned factors, glucosinolates tend to vary quantitatively and qualitatively based on plant part as observed in kale [19], in cabbage [17], and in Arabidopsis thaliana [15].

Analysis of glucosinolates in *Brassica* vegetables has been estimated using HPLC after extraction with boiling water/methanol followed by desulfation of the intact glucosinolates on sephadex-A25 columns [25]. However, desulfation process was seen to be time consuming [26] and some glucosinolates could be insufficiently desulphated at lower concentration of sulphatase [27]. GC-MS methods are often used for detailed analysis [28]. A simplified method of sample extraction and analysis of intact glucosinolates using UPLC-DAD-MS/MS in negative electron-spray ionization (ESI^−^) mode and Multiple Reaction Monitoring (MRM) was employed [29]. In MRM, the precursor/parent ions are selected to make it through the first quadrupole and into the collision cell where they get fragmented. Certain fragment ions (also known as product or daughter ions) are selected to make it through the second quadrupole [30].

Leaves of kimichi cabbage, turnip, Brassica, mibuna, leaf mustard, and cabbage are commonly used for various dishes in many countries. Kimichi cabbage, is a major ingredient in “kimchi”-a widely consumed traditional fermented food in Korea [14]. A number of comparative studies of the profiles of glucosinolates in *Brassica* germplasm collections are available in the literature [1, 13, 20, 31–33]. However, GSLs profiles in large germplasm collections of *Brassica rapa*, *Brassica juncea,* and *Brassica oleracea* is limited except the work of Lee et al. (2014) [14] who identified and quantified ten GSLs of breed varieties of kimichi cabbage collected from the Republic of Korea. Many studies have determined the glucosinolates contents in the seeds [6, 13, 34], and less attention has been given to glucosinolates composition of the leaf of *Brassica,* of which the leaf tissue are usually consumed. Many plant natural products, including glucosinolates, serve as defenses against herbivores [22]. It is important to determine the glucosinolates content in different tissues of the plant to understand their actual defense role a potential herbivore would encounter. In this study, we have identified and quantified the contents of nine GSLs namely, gluconapin, glucobrassicanapin, progoitrin, glucotropaeolin, glucoerucin, gluconasturtiin, glucoberteroin, glucobarbarin, and glucobrassicin in 50 germplasm of *Brassica rapa* subsp. *pekinensis* (kimichi cabbage), *Brassica rapa* subsp. *rapa* (turnip), *Brassica rapa* subsp. *nipposinica* (mibuna), *Brassica juncea* var. *integrifolia* (leaf mustard), and *Brassica oleracea* var. capitate (cabbage) collected form six countries and grown in a uniform agricultural conditions in an attempt to identify differences due to genetic factors. In addition, the accumulation patterns of glucosinolates within and between leaves of kimichi cabbage cultivars were also evaluated.

## Materials and methods

### Reagents and standards

All chemicals and solvents used in extraction and analysis were of analytical grade and purchased from Fisher Scientific Korea Ltd. (Seoul, South Korea) and Sigma-Aldrich (St. Louis, MO, USA). Glucosinolate standards (gluconapin, glucobrassicanapin, progoitrin, glucotropaeolin, glucoerucin, gluconasturtiin, glucoberteroin, glucobarbarin, and glucobrassicin) were purchased from Phytoplan Diehm &Neuberger GmbH (Heidelberg, Germany). All individual GSLs standards were with purity greater than or equal to 97%.

### Plant materials

A total of 50 *Brassica* crops, belonging to *Brassica rapa*, *Brassica juncea*, and *Brassica oleracea* species, and originated from six different countries (China, Ethiopia, Japan, North Korea, South Korea, and Taiwan) was grown at the research farm of the National Agrobiodiversity Center (NAC), Jeonju (35°49′18″ N 127°08′56″ E), Republic of Korea. Seeds of *Brassica* species were sown in plug trays, and seedlings were grown in a greenhouse. Healthy looking seedlings were transplanted to an experimental field. Planting density was 60 x 40 cm. Plant cultural practices were followed in the field as per the recommendation of the Rural Development Administration (RDA), South Korea. Each accession was consisted of 25 plants. Plant growth was maintained using nutrient solution throughout the growing season.

To study the glucosinolate spatial distribution within sections of the leaf of kimichi cabbage and between leaves, two green pigmented (“Hangamssam” and “Alchandul”) and one red pigmented (“Bbalgang 3-ho”) commercial cultivars were selected. The inner, middle, and outer leaves were separated. Each leaf was then dissected into top, middle, bottom, green/red, and white part as required. Sampling positions of kimichi cabbage plant are done as shown in Fig. 1.

**Fig 1.**
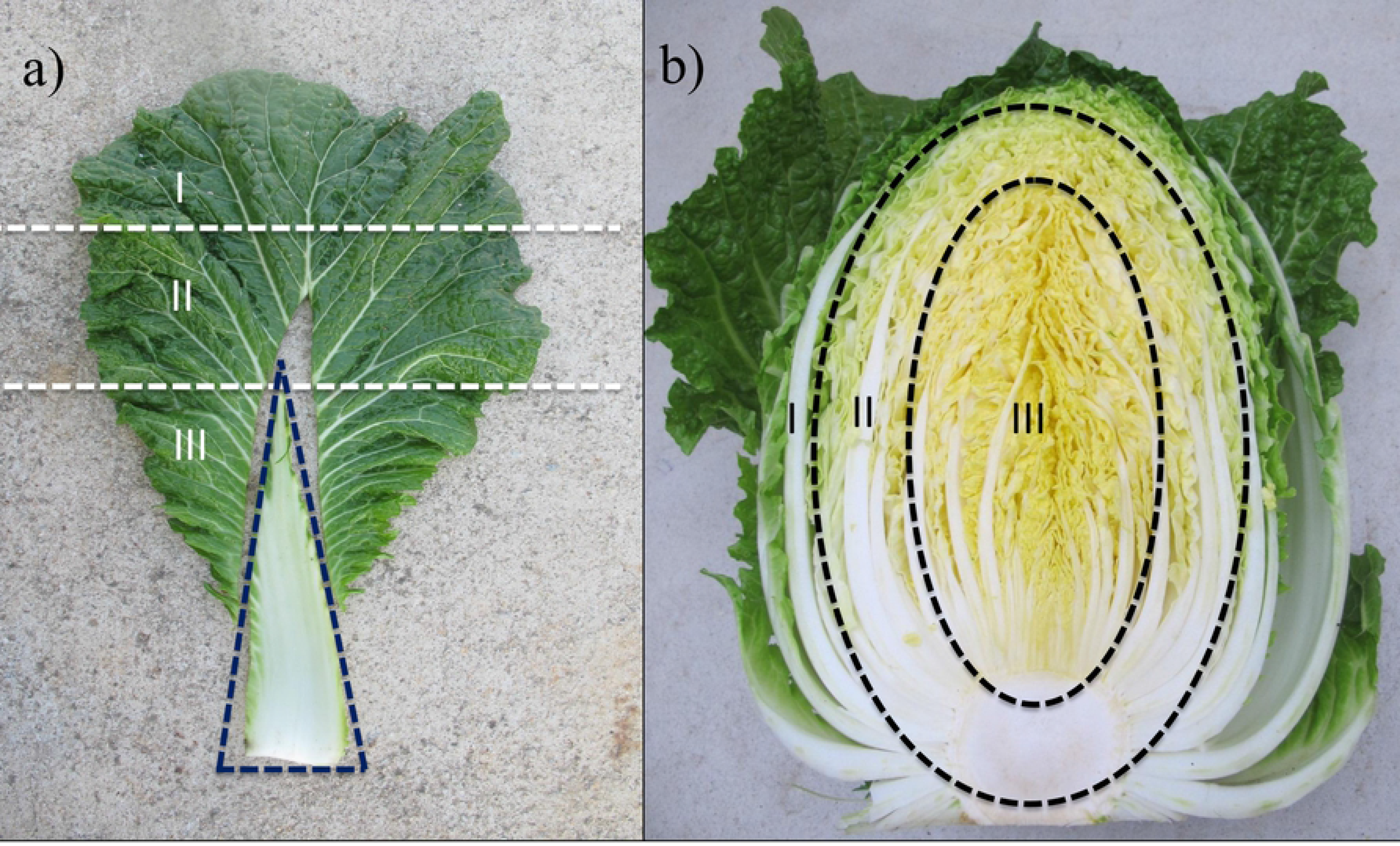
Representative photos of sampling positions of kimichi cabbage based on: **a) Leaf sections: I, III, III refers to the upper, middle and bottom part of the leaf. The white section is indicated in a triangular dashed line. Green/red part was sampled from the whole leaf excluding the white section**. **b) Location of the leaf: I, II, and III refers to the outer (two layers), middle (three layers), and inner (the remaining part) part of the leaves**.

### Sample pre-treatment, extraction and analysis of glucosinolates

Samples were harvested, placed in vinyl freezer bag and kept at −80°C until further processing. The frozen samples were subsequently lyophilized for 48 h using LP500 vacuum freeze-drier (Ilshin biobase Co., Seoul, Korea). The freeze-dried samples were then ground to a fine powder using a mortar and pestle, and held at −80°C until analysis. 0.1 gram of lyophilized sample was mixed with 1 mL of 80% methanol in a 2 mL Eppendorf tube, and sonicated in ultra-sonication bath for 10 min at 30°C. The mixture was centrifuged using VS-180CFi centrifuge (Vision Scientific Co., Daejeon, Korea) (centrifuge conditions set at: 14000 rpm, 4oC, and 10 min). The supernatant was transferred into a vial and glucosinolates were analyzed immediately using UPLC-MS/MS.

The GSLs were analyzed using an Acquity UPLC System (Waters, Milford, MA, USA) coupled to Xevo™ TQ-S system (Waters, MS Technologies, Manchester, UK). Chromatographic separation was carried out using Acquity UPLC BEH C18 (1.7μm, 2.1 × 100mm) column (Waters Corp., Manchester, UK). The flow rate was kept at 0.25 mL/min; the column temperature was maintained at 35°C; and the injection volume was 5 μL. The mobile phase was composed of 0.1% trifluoroacetic acid in water as eluent A and 0.1% trifluoroacetic acid in methanol as eluent B. Elution conditions were as follows: Initial condition set at 100% of A; 0.0 – 1.0 mins, 100 to 95% of A; 1.0 – 4.0 mins, 95 to 0 % A; 4.0 – 4.5 mins, 0 % of A; 4.5 – 5.0 mins, 0 to 100% of A; 5.0 – 10.0 mins, 100% of A. The mass spectrometry instrument was operated in negative ion electrospray ionization (ESI^−^) mode and Multiple reaction Monitoring (MRM). Data acquisition was performed using MassLynx 4.1 software. For MS/MS detection, the ionization source parameters were set as follows: capillary voltage was 3kV; the ion source and the desolvation temperatures were set as 150°C 350°C, respectively. The cone gas (nitrogen) and desolvation gas (also nitrogen) were set at flow rates of 150 and 650 L/h, respectively. Other MRM conditions are presented in Table 1 Glucosinolates were identified by comparing their retention time and MS and MS/MS fragmentation spectra with those of commercial standards. Individual glucosinolates were quantified by MRM, considering one MS/MS transition for each compound. Final concentration of individual glucosinolates was calculated using calibration equations derived from the calibration curves of the corresponding standards. Results are given as μmol kg^−1^ dry weight (DW) sample calculated from LC-peak areas.

**Table 1.** List of identified glucosinolates, retention times, calibration curves, and MRM conditions for quantitation of glucosinolates by negative ion MRM

| Glucosinolates | RT (min) | MRM transition | CV (V) | CID (ev) | Pol. | LOQ (ppb) | Calibration curve parameters |
| --- | --- | --- | --- | --- | --- | --- | --- |
| Gluconapin | 2.26 | 371.89 > 96.0 | 54 | 18 | ES<sup>-</sup> | 1.00 | $Y = 12.376X - 13.4682$ ($r^2 = 0.999$) |
| Glucobrassicinapin | 2.56 | 385.95 > 96.0 | 54 | 17 | ES<sup>-</sup> | 1.00 | $Y = 32.466X - 18.5492$ ($r^2 = 0.994$) |
| Progoitrin | 1.11 | 387.91 > 96.0 | 54 | 18 | ES<sup>-</sup> | 1.00 | $Y = 7.3288 X - 3.38502$ ($r^2 = 0.999$) |
| Glucotropaeolin | 2.58 | 407.97 > 96.0 | 54 | 18 | ES<sup>-</sup> | 1.00 | $Y = 45.1909 X - 47.6141$ ($r^2 = 0.993$) |
| Glucoerucin | 2.61 | 419.95 > 96.0 | 54 | 18 | ES<sup>-</sup> | 1.00 | $Y = 13.135 X - 16.545$ ($r^2 = 0.998$) |
| Gluconasturtiin | 2.91 | 421.97 > 96.0 | 53 | 19 | ES<sup>-</sup> | 1.00 | $Y = 22.8788 X - 21.2328$ ($r^2 = 0.999$) |
| Glucoberteroin | 2.94 | 433.95 > 96.0 | 54 | 18 | ES<sup>-</sup> | 1.00 | $Y = 9.52375 X - 3.64797$ ($r^2 = 0.998$) |
| Glucobarbarin | 2.79 | 437.93 > 96.0 | 52 | 18 | ES<sup>-</sup> | 1.00 | $Y = 9.84022 X - 13.2052$ ($r^2 = 0.996$) |
| Glucobrassicin | 2.68 | 446.95 > 96.0 | 54 | 18 | ES<sup>-</sup> | 1.00 | $Y = 17.5328 X - 19.2757$ ($r^2 = 0.999$) |

CID= Collision Induced Dissociation; LOQ = Limit of Quantification, Pol. = Polarity, CV= Cone Voltage

### Statistical analysis

Results were expressed as mean ± standard deviation (SD) of triplicates. The data were treated with Analysis of Variance (ANOVA) followed by Duncan’s multiple range test (p < 0.05) using the SPSS V. 17.0 statistical program (SPSS Inc., Chicago, USA). The glucosinolate contents in 50 germplasm of *Brassica* were analysed using Principal Component Analysis (PCA). The PCA was performed using PAST (Palaeontological statistics, version 3.06) [35]. Data were visualized using principal components score and loading plots. Each point on the score plot represented an individual sample, and each line on the loading plot represented the contribution of an individual glucosinolate to the score.

## Results

In this study, the GSLs profiles and their concentration were examined in leaves of five commercial varieties and 45 germplasm collections of *Brassica* plants belonging to *B. rapa*, *B. juncea*, and *B. oleracea*. The concentrations of GSLs were also evaluated in various leaf sections and positions of two green (“Hangamssam” and “Alchandul”) and a red (“Bbalgang 3-ho”) pigmented commercial *Brassica* varieties. Five aliphatic (GNA, GBN, PRO, ERU, and BER), three aromatic (TRO, NAS, and BAR) and an indolic (GBC) GSLs were identified. GSLs were examined by negative ionization electrospray LC-MS/MS using MRM mode by monitoring specific transitions originating the characteristic fragment ion at m/z 96 [SO4]^−^. The detection and quantification conditions of the GSLs by LC-MS/MS are presented in Table 1. Information about the germplasm collections and commercial cultivars are presented as a supplementary file (S1 Table). The results of this study, which are to be presented and discussed in detail in the next sections, show the values varied widely among the entire germplasm collections and between different sections and positions of the *Brassica* leaves. Principal Component Analysis (PCA) helps to identify the glucosinolate exhibiting the greatest variance within a population and to determine closely related individual glucosinolates [36]. The data obtained were subjected to PCA to evaluate the glucosinolate difference among the germplasm collections.

### Variation in GSL content between germplasm collections

As can be seen in Table 2, a significant difference in GSL content was observed among the germplasm collections and commercial varieties of *Brassica* plant. The total GSL content ranged from 36.80 (“Alchandul”) to 2383.12 (“Shingatsuna”, IT 135409) μmol kg^−1^ DW with an average value of 951.5 μmol kg^−1^ DW. Aliphatic GSLs were predominant throughout the entire collections which altogether represented from 22.9 to 98.9% (average 79.9%) of the total GSL content, followed by aromatic GSLs (0.2 to 60.4%; average 9.9%). Glucobrassicin, the only indolic GSL detected in our study represented as low as 0.97% and as high as 58.6% with an average of 10.2% of the total glucosinolates. GNA (ranging from 3.8 to 1,589.8μmol kg^−1^ DW) and GBN (ranging from 0.06 to 800.0 μmol kg^−1^ DW) were the most dominant GSLs across the entire collections followed by NAS, GBC, and BER. GNA and GBN represented the greatest proportion (on average 49.8% and 32.3%, respectively) of the total glucosinolates in the leaves of the entire *Brassica* germplasm collection and commercial varieties. NAS, GBC, and BER had represented moderate proportion (7.2%, 5.0%, and 4.7%, respectively) of the total glucosinolates. The least dominant GSLs, TRO, BAR, and PRO contained an average values of 0.30, 0.84, and 1.69 μmol kg^−1^ DW in the entire collections, respectively. GNA and GBN were also documented as the most abundant GSLs in the leaves of kimichi cabbage in previous reports [14, 36]. However, GBN, 4-methoxyglucobrassicin, and progoitrin were the dominant GSLs of the leaves of kimichi cabbage in another study [37]. The identity and quantity of glucosinolates varies considerably between various crops of *Brassica*. For example, the predominant GSLs in broccoli were glucoraphanin, gluconapin, and glucobrassicin, while singrin was found to be the dominant GSL in green cabbage, Brussels sprouts, cabbage, cauliflower, and kale [13, 25].

**Table 2.** Glucosinolates contents (μmol/kg DW) in 50 germplasm accessions of *Brassica* (n=3)

| S/No | GNA | GBN | PRO | TRO | ERU | NAS | BER | BAR | GBC |
| --- | --- | --- | --- | --- | --- | --- | --- | --- | --- |
| 1 | 922.77±0.23uv | 430.77±6.70pqr | 2.65±0.22o | 0.30±0.02lm | 2.35±0.04bcdefg | 83.68±1.40pq | 27.90±0.93jkl | 3.22±0.08qr | 34.91±1.04m |
| 2 | 858.01±21.89rs | 539.65±8.11u | 3.70±0.54qr | 0.24±0.01hij | 1.35±0.12abcd | 154.89±1.49wx | 33.61±2.41klmn | 7.60±0.72s | 28.05±0.41ijk |
| 3 | 828.48±14.53q | 595.11±4.17vw | 4.31±0.50s | 0.24±0.01hij | 2.94±0.20defghi | 117.28±1.94rst | 63.82±4.27q | 3.40±0.13r | 72.11±2.67r |
| 4 | 892.35±12.94t | 557.32±7.80u | 5.14±0.09t | 0.13±0.01ef | 4.86±0.15ij | 163.10±4.28wx | 77.69±2.75r | 0.71±0.03ijk | 34.80±0.27m |
| 5 | 1368.50±14.26x | 216.05±2.18fg | 0.17±0.03abc | 0.66±0.01u | 1.34±0.12abcd | 22.14±0.22defgh | 1.68±0.06ab | 2.90±0.08p | 16.52±0.18de |
| 6 | 757.82±27.74p | 550.05±14.74u | 0.72±0.01defgh | 0.08±0.00bcd | 1.60±0.03abcde | 139.58±4.14v | 40.74±2.11no | 0.78±0.12ijk | 36.19±1.12m |
| 7 | 842.23±40.05qr | 422.46±11.81pq | 0.43±0.04abcdef | 0.11±0.01de | 1.77±0.11abcde | 153.83±5.01wx | 30.12±1.87ijklm | 0.86±0.02jkl | 19.84±0.71efg |
| 8 | 775.31±24.07p | 620.05±13.48w | 0.12±0.01a | 0.13±0.01ef | 0.46±0.01ab | 43.74±1.27jklm | 8.02±0.39abcde | 0.40±0.02cdefg | 35.18±0.79m |
| 9 | 1151.81±10.17w | 270.26±2.79ijk | 0.42±0.04abcdef | 0.18±0.00g | 0.66±0.04abc | 14.81±0.13bcdef | 2.59±0.16ab | 0.88±0.01kl | 22.22±0.20fgh |
| 10 | 598.74±13.78m | 492.80±7.73t | 1.79±0.17mn | 0.26±0.01jk | 1.63±0.16abcde | 119.51±2.19rst | 31.69±3.26ijklm | 1.39±0.03n | 52.52±1.19p |
| 11 | 915.47±20.91u | 371.17±7.96m | 1.04±0.04ghijk | 0.24±0.00hij | 14.72±0.15n | 70.95±0.54o | 60.99±0.74pq | 0.44±0.01defgh | 53.33±0.86p |
| 12 | 866.94±8.93s | 443.92±3.46qrs | 0.21±0.03abc | 0.45±0.02r | 0.22±0.02a | 25.03±0.34fgh | 2.43±0.04ab | 0.31±0.00abcdef | 17.93±0.28ef |
| 13 | 596.00±3.71m | 403.66±2.10nop | 1.14±0.17hijkl | 0.06±0.00ab | 0.74±0.02abc | 112.63±0.57r | 14.91±0.55ef | 0.68±0.04hijk | 33.07±0.37lm |
| 14 | 767.16±8.30p | 250.16±3.54hij | 0.09±0.02a | 0.18±0.00g | 42.96±1.69p | 90.55±1.71q | 173.72±6.55v | 0.57±0.02fghi | 32.93±0.38lm |
| 15 | 722.91±9.40o | 475.35±2.70st | 1.5±0.05lm | 0.34±0.01no | 0.49±0.02ab | 18.31±0.22cdefg | 12.00±0.51cdef | 0.67±0.03hijk | 24.58±0.08ghi |
| 16 | 609.98±8.43m | 552.97±4.18u | 2.09±0.06n | 0.18±0.01g | 1.59±0.12abcde | 87.95±0.99q | 25.73±1.80ijk | 0.54±0.03efghi | 23.45±0.09ghi |
| 17 | 686.80±8.77n | 382.68±2.95mno | 1.00±0.02ghijk | 0.12±0.01e | 7.05±0.33kl | 152.39±0.89w | 33.11±1.8jklmn | 1.10±0.01lm | 29.61±0.33jkl |
| 18 | 820.76±9.14q | 214.97±3.21fg | 0.60±0.02bcdefg | 0.24±0.00hij | 18.18±0.51o | 51.87±0.62lmn | 36.13±1.26lmn | 0.31±0.06abcdef | 41.79±0.47n |
| 19 | 874.87±15.09st | 219.73±2.38fgh | 0.50±0.02abcdef | 0.81±0.02w | 12.54±0.36m | 25.87±0.30fgh | 34.21±0.91lmn | 0.29±0.03abcde | 21.24±0.29efgh |
| 20 | 384.29±6.14k | 458.45±97.47rs | 3.54±0.39q | 0.41±0.05q | 8.17±0.57l | 164.25±33.26x | 152.10±7.00u | 1.01±0.1lm | 101.13±12.02v |
| 21 | 1589.80±2.92y | 567.51±3.28uv | 1.44±0.05klm | 0.19±0.01g | 7.29±0.17kl | 131.75±1.23uv | 25.17±0.76hij | 0.44±0.05defgh | 59.53±0.67q |
| 22 | 210.67±2.36h | 290.45±2.76kl | 6.46±0.66v | 0.72±0.01v | 13.47±0.18mn | 124.72±2.74stu | 139.69±1.26t | 1.01±0.11lm | 285.68±6.23x |
| 23 | 940.19±15.88v | 65.22±0.50cd | 0.04±0.01a | 0.41±0.01q | 0.12±0.01a | 1.41±0.02a | 0.07±0.01a | 0.05±0.01a | 9.43±0.06abc |
| 24 | 462.58±15.94l | 454.74±7.95qrs | 0.28±0.00abcd | 0.34±0.01no | 1.09±0.04abcd | 74.81±1.58op | 17.98±1.16fgh | 0.35±0.07bcdef | 27.38±0.90ij |
| 25 | 229.76±9.34h | 408.72±12.94op | 2.57±0.17o | 0.60±0.02st | 4.20±0.32ghi | 152.26±4.98w | 115.53±8.08s | 1.14±0.07m | 87.96±2.41t |
| 26 | 292.95±1.79j | 373.93±7.24mn | 1.77±0.09mn | 0.28±0.02kl | 12.49±0.56m | 84.72±2.71pq | 216.00±11.01w | 0.45±0.08defgh | 47.94±1.22o |
| 27 | 180.12±3.30g | 429.44±4.61pqr | 2.09±0.07n | 0.82±0.00w | 4.60±0.26hij | 164.54±3.17x | 84.08±5.69r | 2.23±0.05o | 69.80±1.40r |
| 28 | 1132.99±9.19w | 800.03±9.00x | 1.23±0.06ijkl | 0.10±0.02cde | 0.53±0.05ab | 126.91±1.79tu | 30.28±0.99ijklm | 0.62±0.03ghij | 25.03±1.73hij |
| 29 | 683.37±11.91n | 193.19±6.33f | 0.13±0.03ab | 0.05±0.00ab | 1.47±0.06abcd | 9.60±0.35abc | 3.97±0.25abc | 0.12±0.00ab | 9.52±0.14abc |
| 30 | 258.39±13.36i | 457.99±12.33rs | 7.22±0.72w | 0.35±0.01nop | 3.63±0.28efghi | 32.72±1.22hij | 55.93±1.94p | 0.29±0.02abcde | 13.05±0.36cd |
| 31 | 99.27±0.53e | 314.34±4.23l | 5.68±0.18u | 0.25±0.00ijk | 1.18±0.07abcd | 72.95±0.62o | 36.08±0.63lmn | 0.21±0.03abcd | 96.64±0.86uv |
| 32 | 59.57±1.81cd | 214.42±5.45fg | 3.07±0.05p | 0.30±0.00lm | 3.63±0.19efghi | 45.33±0.61klm | 57.07±2.41pq | 0.14±0.01abc | 97.81±2.13uv |
| 33 | 53.52±2.15cd | 146.53±4.03e | 2.51±0.11o | 1.12±0.05x | 2.73±0.21cdefgh | 57.29±1.61n | 47.13±4.41o | 0.27±0.01abcd | 61.98±2.85q |
| 34 | 125.99±0.40f | 243.15±1.94ghi | 0.88±0.11fghij | 0.38±0.01pq | 0.31±0.00ab | 37.75±0.25ijk | 12.39±0.59def | 0.17±0.01abc | 78.37±0.39s |
| 35 | 233.55±2.21h | 280.20±0.85jk | 1.02±0.06ghijk | 0.21±0.01h | 0.60±0.04abc | 53.42±0.17mn | 12.11±0.59cdef | 0.29±0.00abcde | 34.21±0.43lm |
| 36 | 130.69±3.18f | 212.84±2.55fg | 1.00±0.02ghijk | 0.28±0.01kl | 6.23±0.09jk | 43.08±0.55jklm | 54.46±0.17p | 0.18±0.03abcd | 96.00±1.35u |
| 37 | 15.24±0.70a | 28.17±0.55ab | 0.15±0.01ab | 0.37±0.01op | 0.55±0.05ab | 41.62±0.55jkl | 8.50±0.86bcde | 0.21±0.03abcd | 134.02±0.39w |
| 38 | 40.88±0.65bc | 78.44±0.92cd | 0.50±0.15abcdef | 0.15±0.00fg | 1.92±0.06abcdef | 15.06±0.15bcdef | 17.09±0.38fg | 0.10±0.02ab | 54.15±0.16p |
| 39 | 27.20±1.60ab | 85.97±2.79d | 0.80±0.07efghi | 0.23±0.00hi | 0.63±0.08abc | 27.62±1.15ghi | 15.81±2.47ef | 0.09±0.00ab | 43.91±1.20no |
| 40 | 102.91±2.16e | 353.98±5.67m | 3.45±0.15pq | 0.08±0.00abc | 0.61±0.05abc | 43.70±0.47jklm | 24.17±2.06ghi | 0.19±0.01abcd | 6.46±0.05a |
| 41 | 75.65±3.25d | 146.77±4.62e | 1.30±0.07jkl | 0.08±0.01bcd | 3.87±0.11fghi | 42.88±0.99jklm | 36.66±1.45mn | 0.15±0.01abc | 88.98±1.95t |
| 42 | 396.05±6.22k | 559.56±3.53u | 3.43±0.07pq | 0.09±0.00cd | 1.24±0.01abcd | 114.12±2.32rs | 44.89±1.41o | 3.13±0.03q | 43.08±0.84n |
| 43 | 20.59±0.82ab | 90.50±2.29d | 0.77±0.09efgh | 0.17±0.01g | 0.88±0.05abcd | 13.52±0.26bcde | 36.78±0.99mn | 0.21±0.03abcd | 6.24±0.10a |
| 44 | 3.88±0.18a | 14.43±0.63a | 0.29±0.02abcd | 0.21±0.01h | 1.26±0.09abcd | 5.62±0.21ab | 5.97±0.51abcd | 0.05±0.01a | 5.09±0.19a |
| 45 | 7.85±0.16a | 13.62±0.27a | 0.28±0.02abcd | 0.07±0.00abc | 0.67±0.09abc | 5.44±0.06ab | 8.89±1.80bcde | 0.10±0.00ab | 8.64±0.07abc |
| 46 | 7.48±0.12a | 17.78±0.17a | 0.12±0.02a | 0.05±0.00a | 0.41±0.01ab | 6.57±0.06ab | 5.20±0.14abcd | 0.08±0.01ab | 11.38±0.04bc |
| 47 | 27.45±1.52ab | 0.06±0.01a | 0.41±0.03abcde | 0.44±0.02r | 0.12±0.02a | 7.97±0.38abc | 0.01±0.00a | 0.08±0.01ab | 32.21±0.74klm |
| 48 | 6.26±0.08a | 5.68±0.01a | 0.64±0.02cdefg | 0.33±0.01mn | 1.00±0.03abcd | 12.23±0.36abcd | 8.73±0.07bcde | 0.15±0.01abc | 18.55±0.44ef |
| 49 | 9.73±0.33a | 0.15±0.01a | 0.05±0.00a | 0.58±0.00s | 0.01±0.00a | 24.47±1.25efgh | 0.01±0.00a | 0.33±0.03abcdef | 6.66±0.28ab |
| 50 | 44.43±0.63bc | 53.25±1.38bc | 4.00±0.05rs | 0.63±0.01t | 83.06±5.87q | 56.94±0.11n | 247.65±14.4x | 0.87±0.05jkl | 100.97±1.31v |

GNA = Gluconapin; GBN = Glucobrassicinapin; PRO= Progoitrin; TRO = Glucotropaeolin; ERU = Glucoerucin; NAS = Gluconasturtiin; BER = Glucobetteroin; BAR = Glucobarbarin; GBC = Glucobrassicin

The results of PCA are indicated by the principal components score and loading plots. The PCA of glucosinolates data yielded three principal components with eigenvalue ≥1, accounting 70.33% of the total variance across the entire data set. The first, second, and the third principal components (PCs) contributed 32.17, 25.6, and 12.18 % of the total variance, respectively. The loadings, eigenvalues, and percentage of variance for all principal components (PCs) yielded are attached in a supplementary file (S2 Table). Scores and loading plots of PC1 and PC2 of the PCA results obtained from glucosinolate content of 50 *Brassica* germplasm collections are presented in Fig 2. The loadings of glucosinolates (represented by green lines) show the extent of each glucosinolate concentration contributed to the principal components. All the glucosinolates were positively correlated with PC1 while only GNA, GBN, BAR, and NAS had a positive correlation with PC2. NAS was the predominant glucosinolate in PC1 while the aliphatic glucosinolates GNA and GBN had a major contribution to PC2. Three kimichi cabbage samples (2, 22 and 50) (see S1 Table for more), the first one located at the top right hand quadrant and others at the lower right hand quadrant of the PCA plot, were well distinguished from other samples. The location of these materials in the score plot could be described by their significantly higher content of BAR, GBC and ERU respectively.

**Fig 2.**
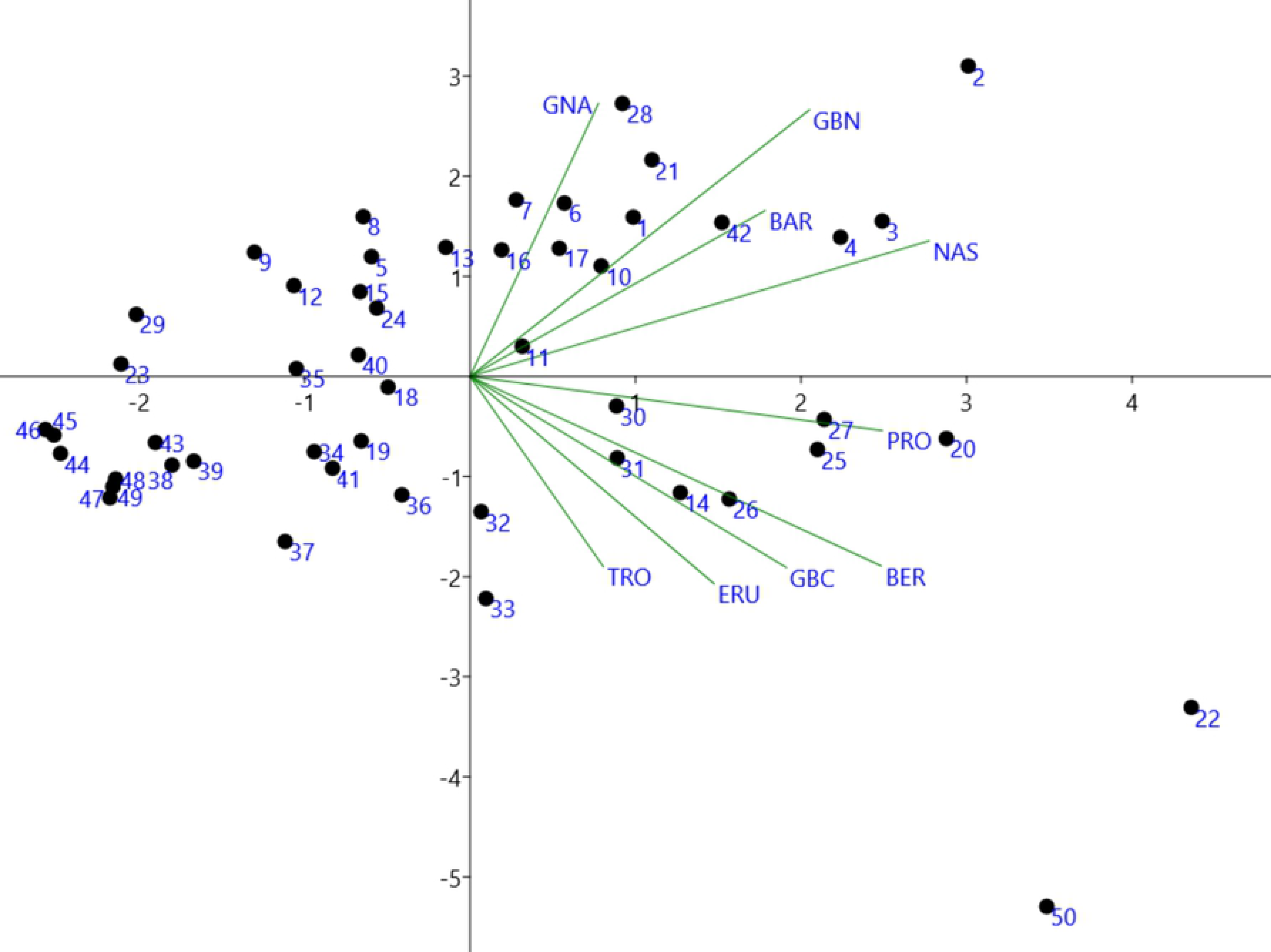
Principal Component Analysis (PCA) plot of the scores (indicated by dotes) and loadings (indicated by lines) of the 50 *Brassica* plants based on the first and second principal components. The numbers 1-50 corresponds to the S/No in Table 2 and S1 Table. GNA = Gluconapin; GBN = Glucobrassicanapin; PRO= Progoitrin; TRO = Glucotropaeolin; ERU = Glucoerucin; NAS = Gluconasturtiin; BER = Glucoberteroin; BAR = Glucobarbarin; GBC = Glucobrassicin.

### Intra-and inter-leaf distribution of glucosinolates in kimichi cabbage

The leaves of three cultivars of green/ red pigmented kimichi cabbage namely, “Hangamssam” (green), “Alchandul” (green), and “Bbalgang 3-ho” (red) segregated based on their position in the whole leaf part as inner, middle, and outer parts and each leaf was further portioned into different sections (top, middle, bottom, green/red, and white). The GSLs content in kimichi cabbage significantly varied based on leaf section, position, and color. The GSLs content in different leaf sections/positions of three cultivars of kimichi cabbage are presented in Table 3. The white sections of the leaf contained higher total sum of glucosinolates (1.16 to 24.28-fold higher) than the green/red section except in the outer leaf of “Bbalgang 3-ho” where the red section contained 2.8-fold greater total sum of GSLs concentration than the white section. The trend in total GSLs content in different sections of the leaf (top, middle, and bottom) was not strictly consistent. However, in most cases higher GSLs content were observed at the proximal half of the leaves. In regard to the position of the leaf (outer, middle, and inner) in the whole plant, the average content of total GSLs of the kimichi cabbage in the middle position were 1.8-, 2.2-, and 3.9-fold larger than in the outer positions of “Hangamssam”, “Alchandul”, and “Bbalgang 3-ho”, respectively. The content of total glucosinolates evaluated in the inner position of “Alchandul” and “Bbalgang 3-ho” showed 2.8- and 1.2-fold higher than the outer position, respectively. However, unlike “Alchandul”, less total GSLs content was recorded in the red pigmented “Bbalgang 3-ho” in the inner compared to the middle leaf. In general, the younger (inner) leaves were found to contain higher concentrations of glucosinolates compared to older (outer) leaves. In earlier study, a similar trend was observed in *Brassica oleracea* var. *capitate* where the inner positions contained 1.1- to 1.8-fold greater GSLs concentrations than the outer positions [38]. In another study form *Raphanus sativus*, younger leaves were found to contain higher glucosinolate concentrations [39].

**Table 3.**
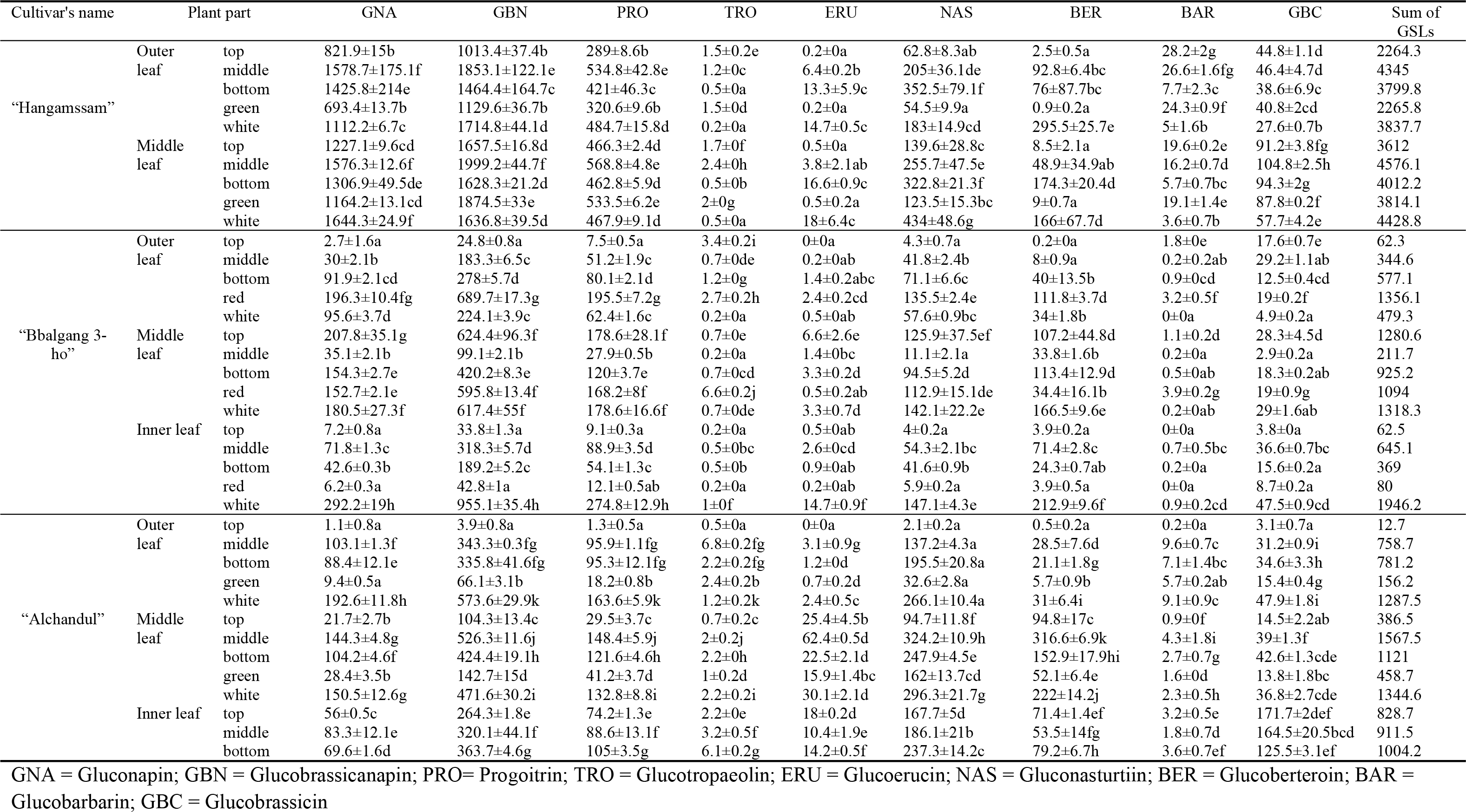
Glucosinolate concentration in different leaf sections of three cultivars of kimichi cabbage (μmol/kg DW)

## Discussions

The data in this study, generated from evaluations conducted from *Brassica* germplasm collections and commercial cultivars originated from various countries and grown in the same location and season, revealed that a wide variety in the level of glucosinolates among genotypes, leaf position/section, and leaf color was observed. The difference observed in GSLs profile is both qualitative and quantitative. This could determine their level of nutritional and health promoting properties and supports the feasibility of developing germplasm with enhanced level of glucosinolates through genetic manipulation. Previous studies have shown that the temperature [19], amount of rainfall [40], radiation [41, 42], plant part examined [1], phenological stage of growth [15, 19], and level of insect damage [19, 43] have affected the level of glucosinolates. The degradation products (pent-4-enyl-isothiocyante and 5-(methylsulfinyl) pentyl isothiocyante) of the dominant GSLs found in this study, gluconapin and glucobrassicanapin, respectively, found to inhibit a wide range of bacteria and fungi indicating their promising antimicrobial potential [6]. The indolic glucobrassicin, the aliphatic glucoberteroin, and the aromatic gluconastrutiin were found in appreciable amount in our study, suggesting that the germplasm collections could be beneficial in terms of the health related activities of these compounds. The derivatives of GBC, indole-3-carbinol (I3C) and 3,3’-diindolylmethane (DIM), have been reported to induce apoptosis to suppress the growth of human prostate cancer cells *in vitro* [44], while glucoberteroin showed to strongly suppress protein glycation and carbonylation which in turn causes aging of the skin [45]. The hydrolysis product of NAS, phenethyl isothiocyanates (PEITC), shown to induce apoptosis in HeLa cells in a time- and dose-dependent manner [46]. In another study, PEITC was reported to create an oxidative cellular environment that induces DNA damage and *GADD153* gene activation, which in turn helps trigger apoptosis [47].

The enhancement of glucosinolate concentration upon plant damage [43], have long indicated that glucosinolates are plant defense chemicals where mostly their defensive properties are attributed to the toxicity and deterrence nature of their degradation products [9]. However, there are also cases that glucosinolates mediated by their volatile hydrolysis products could serve to attract adapted herbivores that often use glucosinolates as cues for feeding or oviposition [9]. The spatial distribution of glucosinolates in different section of a single leaf and/or location of the leaf in the whole plant could be partly important to explain the patterns of herbivory. Studies devoted to glucosinolate spatial patterns within leaves of kimichi cabbage are elusive. The proximal half of leaves of *Raphanus sativus* contained higher mean concentration of glucosinolates compared to the distal halves of leaves [39]. Shroff et al. (2008) [48] studied the spatial distribution (midvein, inner lamina, and outer lamina) of glucosinolates in leaves of *Arabidopsis thaliana* and tried to relate the distribution to the pattern of herbivory caused by larvae of the lepidopteran, *Helicoverpa armigera.* These authors found out that the glucosinolate abundance in the inner vs. the peripheral part of the leaf affected insect feeding preference and anti-herbivore defenses. As stated in previous section and Table 3, the white part (midvein) of kimichi cabbage contained relatively higher glucosinolates compared to the green or red. This is consistent with *Arabidopsis thaliana* leaves where the midvein part exhibited the greatest concentration compared to the other sections of the leaf [48]. This could be due to certain biosynthetic enzymes are distributed exclusively to vascular bundles [49], resulting greater synthesis and storage of glucosinolates in the midvein (white part) of the leaf of kimichi cabbage. It could also be related to ecological significance [50], the midvein being critical to the function of the leaf as the transport of water and nutrients are carried through it. The greater concentration of glucosinolate in the white part of kimichi cabbage in our study corroborates the idea of storage of glucosinolates being associated with the vascular system. The higher content of glucosinolates in the inner (younger) leaves compared to the outer (older) leaves in this study is also in agreement with the predictions of optimal defense theory: young leaves are more valuable as they have a higher future photosynthetic potential needs higher degree of protection from damage [51]. In addition, glucosinolate concentration could tend to decrease in outer leaves due to dilution of glucosinolates as the leaf expands [51].

## Conclusions

Nine glucosinolates were identified and quantified in *Brassica* germplasm collections and commercial varieties using UPLC-MS/MS method in Multiple Reaction Monitoring scan mode. Remarkable differences in total and individual glucosinolates were observed among different samples. The variation in glucosinolate level suggests that the potential health benefits of *Brassica* plants could depend on the type of accession used. Glucosinolates and their degradation products are known for their chemopreventive properties, and the wide variability in glucosinolates among the germplasm collections in this study offer important information base for enhancing the level of glucosinolates in *Brassica* plants through breeding thereby enhancing their anti-cancer properties. In addition, the development of *Brassica* plants with specific GSL profiles of specific functions will allow for meaningful recommendations of dietary intake of *Brass*ica vegetables in respect to other biological activity. The PCA in this study allowed easy visualisation of the data; three kimichi cabbage samples (2, 22, and 50) were separated from the others in The PCA plot. The inter- and intra-leaf variations of GSLs were examined in three kimichi cabbage varieties. The GSLs content varied significantly among leaves in different positions of the plant (outer, middle, and inner) and sections within leaves (top, middle, bottom, green/red, and white). Higher GLS contents were observed the proximal half and white section of the leaves and inner positions (younger leaves) in most of the samples. Studying how the variability of GSLs content and composition is reflected within and between leaves would widen the present understanding of the accumulation pattern of GSLs in leaves of *Brassica* plants and provide information on the nature of plant defenses towards perceived danger. To better understand the extent and pattern of anti-herbivore glucosinolate defenses, further investigation of glucosinolate distribution in relation to the pattern of herbivory by insects is recommended.

## Acknowledgements

This research was carried out with the support of “Research Program for Agricultural Science & Technology Development (Project NO. PJ01425501)”, National Institute of Agricultural Sciences, Rural Development Administration, Republic of Korea.

## Supporting information

S1 Fig. Representative MRM profiles of intact glucosinolates corresponding to Brassica germplasm

S1 Table. Accession number, scientific name, common name and origin of 50 germplasm accessions of *Brassica* genus

S2 Table. Loadings, eigenvalues, and percentage of variance for the principal components (PCs) data from germplasm collections

